# Stimulating the regenerative capacity of the human retina with proneural transcription factors in 3D cultures

**DOI:** 10.1101/2024.09.24.614778

**Authors:** Juliette Wohlschlegel, Faith Kierney, Kayla L. Arakelian, Guillaume Luxardi, Naran Suvarnpradip, Dawn Hoffer, Fred Rieke, Ala Moshiri, Thomas A. Reh

**Author notes:** Correspondence should be addressed to T.A.R.

## Abstract

Retinal diseases often lead to degeneration of specific retinal cell types with currently limited therapeutic options to replace the lost neurons. Previous studies have reported that overexpression of ASCL1 or combinations of proneural factors in Müller glia (MG) induces regeneration of functional neurons in the adult mouse retina. Recently, we applied the same strategy in dissociated cultures of fetal human MG and although we stimulated neurogenesis from MG, our effect in 2D cultures was modest and our analysis of newborn neurons was limited. In this study, we aimed to improve our MG reprogramming strategy in a more intact retinal environment. For this purpose, we used an in vitro culture system of human fetal retinal tissue and adult human postmortem retina. To stimulate reprogramming, we used lentiviral vectors to deliver constructs with a glial-specific promoter (HES1) driving ASCL1 alone or in combination with additional developmental transcription factors such as ATOH1 and NEUROD1. Combining IHC, scRNA-seq and electrophysiology, we show for the first time that human MG can generate new neurons even in adults. This work constitutes a key step towards a future clinical application of this regenerative medicine approach for retinal degenerative disorders.

## Introduction

In the mammalian central nervous system (CNS), any loss of neurons, whether accidental, such as through injury, or resulting from the process of aging and disease, is an irreversible process; the retina is no exception to the rule. Retinal degeneration is one of the leading causes of blindness, and retinal diseases like age-related macular degeneration impose a great personal as well as an economic burden worldwide. While new therapeutic strategies and technologies are constantly emerging to delay the progression of retinal diseases, restoration of visual function after cellular degeneration has occurred is still not possible. In the field of regenerative medicine, cell transplant and regeneration are two complementary approaches used to replace the damaged cells and ultimately restore visual function(1–3). Regenerative strategies offer the advantage of utilizing endogenous cells, thereby eliminating some of the complexities of cell transplantation, including abrogating the need for immunosuppressant therapies.

Several studies in the last few years demonstrated that Müller glia (MG), the principal retinal glial cells, can be reprogrammed to generate neurons in the adult mouse retina(4– 10). Inspired by the regenerative capacity of non-mammalian vertebrate retina(11–13), this strategy relies on overexpressing the proneuronal transcription factor (TF) Ascl1 in MG (5,6,14). With this strategy, newly generated neurons are integrated into the existing circuit, and molecularly and functionally resemble inner retinal neurons, primarily bipolar cells(4–6). Additional studies have made significant progress boosting the efficiency of MG reprogramming and expanding the cell fate diversity of MG derived neurons in the mouse retina(8,15–17). For instance, the combination of the developmental TFs Islet1 and Pou4f2, along with Ascl1 can reprogram MG into RGC-like cells(7). This work supports the feasibility of restoring retinal cell types in the adult mammalian retina; however, these studies relied on Cre dependent transgenic mouse models, which cannot be directly translated to humans.

In a recent study, we demonstrated that dissociated cultures of human fetal or organoid-derived MG can be reprogrammed into neurons using a lentivirus expressing ASCL1(18). Consistent with previous studies in mice, most of the MG expressing ASCL1 are converted to neurogenic precursors and only a subpopulation generated neurons(18). Interestingly, human MG-derived neurons resemble immature amacrine-like, and RGC-like cells, as opposed to mouse MG-derived neurons that more closely resembled bipolar cells. Although, these results present a proof of principle that human MG can also be reprogrammed into neurons, those neurons do not fully differentiate in dissociated cultures, likely due to a lack of environmental cues.

In this study we aimed to reprogram human MG in a more intact and physiologically relevant retinal environment. For this purpose, we used explant cultures of fetal and adult postmortem retinal tissue. We further designed a new lentiviral strategy that uses the HES1 promoter (19,20) to precisely drive expression of reprogramming TFs in MG, to stimulate neurogenesis in these cells. One major advantage of this approach is the maintenance of the expression of ASCL1, even as cells reprogrammed from glia to progenitor cells, since HES1 is also expressed in progenitor cells during fetal development. Using this approach, we now demonstrate that neural regeneration can be stimulated in both fetal and adult human retinas. This work provides a crucial step towards translation of in vivo reprogramming as a therapy for retinal diseases.

## Results

### HES1 promoter in a lentiviral construct drives expression specifically in human MG. (Fig 1) (Fig S1)

To identify potential promoters that could be used for stimulating reprogramming in MG, we examined Multiome data from a D59 human fetal retina to identity promoters that drive gene expression in both MG and retinal progenitors (18). Our rationale was that these promoters would potentially be active both in the target glial population and the glial-derived progenitors as they are reprogrammed. We identified HES1 as an ideal candidate, since this gene is expressed both in MG and in progenitors in the snRNA-seq data **(Fig 1A)** and the snATAC-seq results show accessibility in both cell types in the region corresponding to the human HES1 promoter **(Fig 1A)**. After confirming the expression of HES1 in human MG **(Fig S1A)**, we then designed and generated lentiviral constructs with the HES1 promoter driving the expression of ASCL1 followed by a fluorescent protein (GFP). We also generated a control vector, with HES1 promoter driving GFP. Retinospheres from different samples and at different ages were infected with the constructs (HES1-GFP and HES1-ASCL1-GFP). To ensure reproducibility, each retinosphere from the same condition was infected individually and received the same cocktail of virus. In addition, only retinospheres older than 200 days were used for the study since we previously demonstrated that MG are well developed, and few progenitors remain by that timepoint (18).

**Figure 1:**
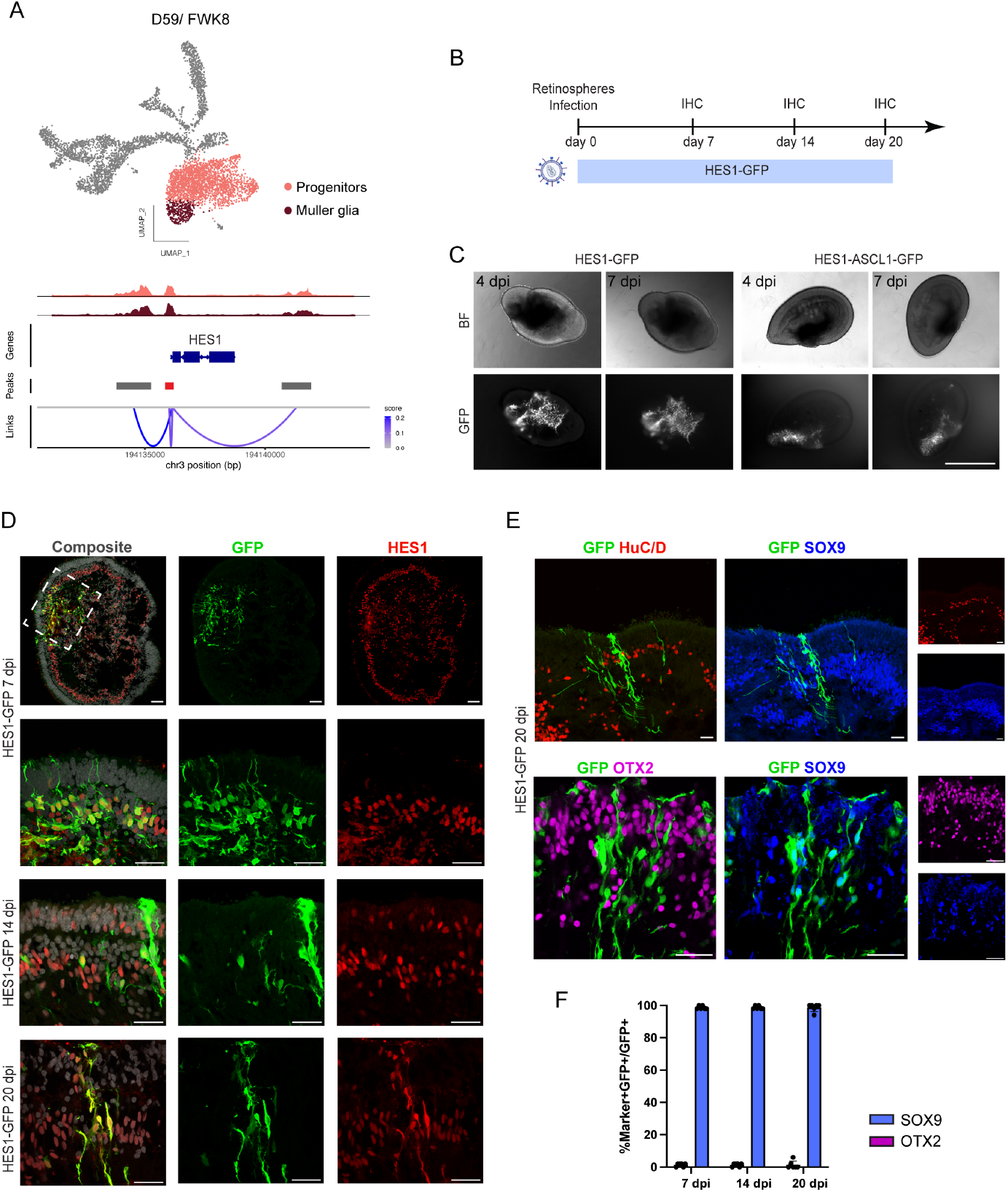
HES1 promoter displays a glial-restricted activity in retinospheres. (A) Multiome data from a D59 fetal human retinal sample. Top: UMAP plot from snRNAseq data organized by cell type (orange; retinal progenitors and dark red: Muller glia). Bottom: coverage plot nearby HES1 gene showing peak to gene analysis, data obtained from the snATAC-seq data. The red sequence represents the presumed sequence of the HES1 promoter. (B) Schematic of the experimental timeline to test the specificity of the HES1 promoter. (C) Retinospheres infected with the different lentiviral constructs at 4- and 7-days post infection. Top panels show the brightfield (BF) images and lower panels show the GFP reporter expression. Scale bar = 500 *µ*m. (D) HES1 expression co-localized with GFP+ cells at 7-, 14-and 20 days post infection with the HES1-GFP virus. DAPI (grey), GFP (green) and HES1 (red). Scale bar = 50 *µ*m (top row); Scale bar = 30 *µ*m (lower panels). (E) Representative images showing that GFP+ cells do not colocalized with the HuC/D (red) nor OTX2 (magenta) neural markers 20 days post infection with the HES1-GFP virus. Scale bar = 30 *µ*m. (F) Quantification showing a residual number of GFP/OTX2+ cells after HES1-GFP infection, (6 retinospheres per timepoint, 7 dpi [2 x D245, 4 x D343], 14 dpi [2 x D209, 4 x D255], 20 dpi [1 x D209, 2 x D255, 2 x D301]).

Retinospheres were infected with the HES1GFP or HES1-ASCL1-GFP viral vectors, fixed at different at time points (7-, 14- and 20-days post infection (dpi)) and later sectioned and immu-nolabelled with different neuronal and glial markers **(Fig 1B)**. By 4 dpi, GFP was already detectable in both conditions **(Fig 1C)**. To test the glial specificity of the HES1 promoter, we then focused on the HES1-GFP condition. At 7 dpi, immunohistochemistry (IHC) analysis revealed that GFP+ cells co-localized with the HES1 glial marker **(Fig 1D)** and that the cell type specificity was maintained at 14- and 20-dpi **(Fig 1D)**. Furthermore, at 20 dpi, GFP+ cells expressed the SOX9 glial marker and did not express OTX2 or HuC/D, two neuronal markers **(Fig 1E)**. Quantification of the GFP+ cells further validated this result showing that the number of GFP/OTX2+ neurons was minimal and constant over time (6 retinospheres per time point, average of 1.33 % at 7-, 14- and 20-dpi) **(Fig 1F)**. These results demonstrated the HES1 promoter exhibits a glial-restricted activity in retinospheres.

### Efficient reprogramming of human MG in 3D retinospheres with the HES1-ASCL1-GFP viral vector. (Fig 2) (Fig S2)

To detect the expression of ASCL1, sections were immunolabelled with a MASH1 (ASCL1) antibody at 7 dpi. In the HES1-ASCL1 infected condition, we observed a robust expression of ASCL1 that was absent in the control condition (HES1-GFP) **(Fig S2A)**. To determine whether the expression of a proneural TF would change the glial specificity of the HES1 promoter, a phenomenon that has confounded previous studies on reprogramming glia into neurons (21–23), retinospheres were infected with the ASCL1 construct and fixed at 2 dpi, when the first GFP+ cells were detectable. We reasoned that 2 days would not be sufficient time for MG to be reprogramed to neurons, so any GFP expression in neurons would indicate a change in the glial specificity of the HES1 promoter. We found that after 2 dpi, GFP+ cells were primarily co-labeled with SOX9, and did not express the OTX2 neuronal marker (%SOX9+GFP+/GFP+, 97.6%) **(Fig S2B)**. Furthermore, we found ASCL1 expression in SOX9/GFP+ cells **(Fig S2C)**. We therefore conclude that during the first two days following the infection, the addition of the ASCL1 to the lentiviral construct did not modify the specificity of the promoter: we saw no evidence of GFP expression in neurons.

**Figure 2:**
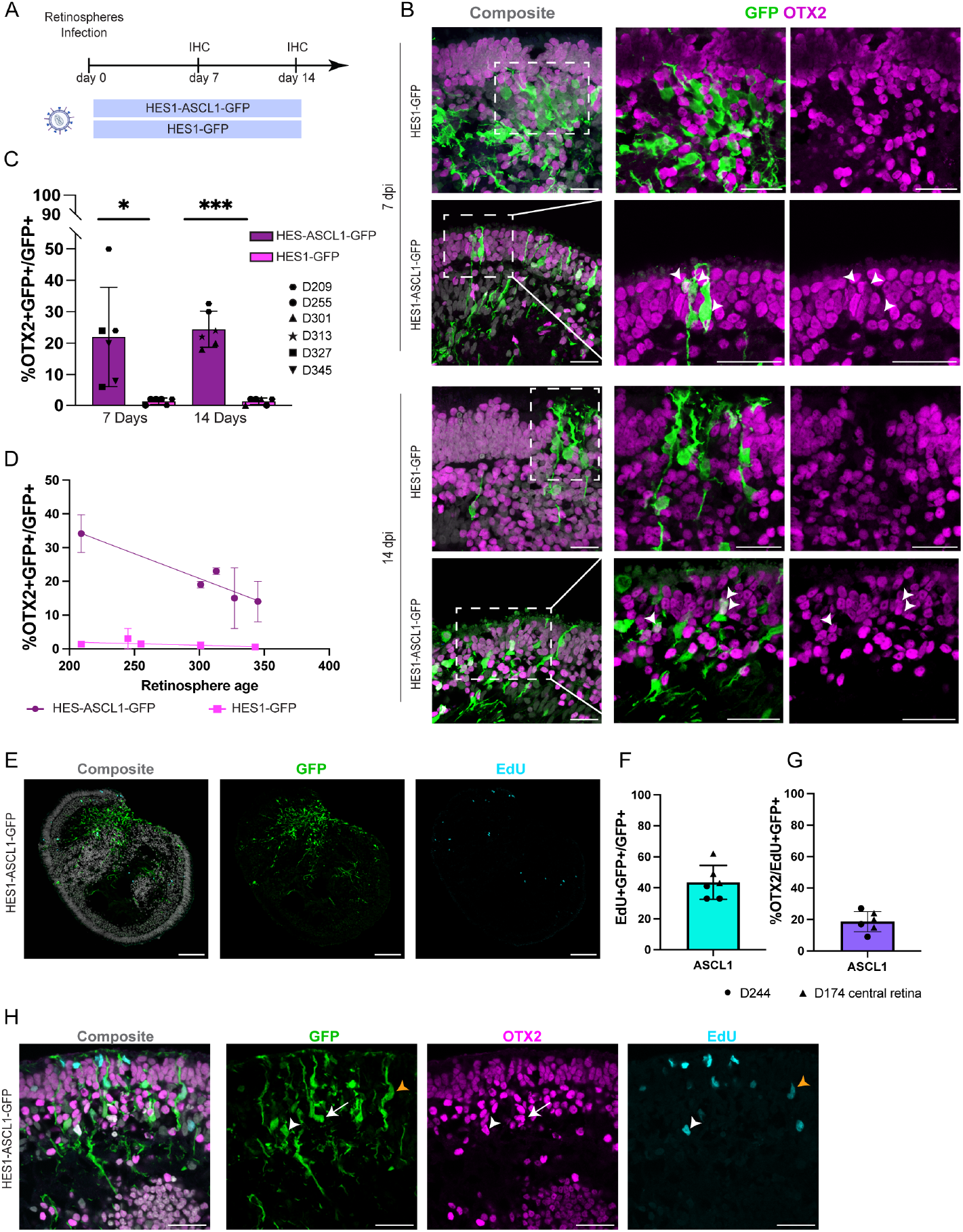
Expression of ASCL1 via the HES1 promoter enables MG reprogramming into neurons in retinospheres. (A) Schematic of the workflow to test MG reprograming with the HES1-ASCL1-GFP lentivirus. (B) Representative images showing MG-derived neurons expressing OTX2 (magenta) after ASCL1 expression at 7-and 14-days post infection. Arrowheads show co-labeled GFP/OTX2+ cells. DAPI (grey). On the left side of the panel, Scale bar = 25 *µ*m. and Scale bar = 30 *µ*m on the right side of the panel. (C) Quantification of the percent of GFP+ MG-derived neurons that express OTX2, 7-and 14-days post infection. Each shape represents a retinosphere. Ages of the retinospheres are indicated on the right side of the figure. Significance was determined by a Welch’s t-test (*p < 0.05, ***p < 0.001). (D) Reprogramming efficiency decreases with the ages of the retinospheres. Each shape (pink dots for HES1-ASCL1 and green squares for HES1-GFP) represents all the combined data for this specific age. Data are represented as mean ± SEM. (E) Representative images showing MG proliferation after ASCL1 overexpression, GFP (green), EdU (cyan) and DAPI (grey). Scale bar = 100 *µ*m. (F) Quantification of GFP+ cells co-labelled with EdU after ASCL1 overexpression. (G) Quantification of MG-derived new-born OTX2+ cells after ASCL1 overexpression. Each shape represents a retinospheres. (H) Confocal images showing MG to neurons conversion. White arrows show GFP/OTX2+ cells, orange arrowheads indicate GFP/EdU+ cells, and white arrowheads show GFP/OTX2/EDU+ cells. Scale bar = 30 *µ*m.

To determine whether longer expression of the proneural TF would induce neurogenesis in the MG, we infected retinospheres as above and then incubated them for 7 or 14 days **(Fig 2A)**. In addition, EdU (5-ethynyl-2’-deoxyuridine) was continuously added to the medium to track the generation of new cells. We found that in the HES1-ASCL1-GFP condition, GFP+ cells acquired a more pronounced neuronal morphology, with a smaller and rounder nucleus, and tended to migrate to the outer nuclear layer (ONL) **(Fig 2B)**. While GFP+ cells from the control condition were all positive for the MG marker SOX9 and negative for the neuronal marker OTX2, 22% and 24% of the GFP+ MG-derived cells from the 7 and 14 dpi treatment groups, respectively, expressed OTX2 **(Fig 2C)**. Other neuronal markers such as HuC/D (expressed by RGCs and amacrine cells) were also tested; however, we did not observe any GFP/HuC/D+ cells (data not shown). Of note, we noticed a decrease in reprogramming efficiency over time as the retinospheres age: approximately 35% of the GFP+ cells generated OTX2+ neurons from retinospheres infected at D209 whereas this number was decreased to only 26% at D300 **(Fig 2D)**.

We then calculated the percentage of infected cells that were EdU+ and found that almost half of the GFP+ cells were EdU+ (43.5% of %EdU+GFP+/GFP+) **(Fig 2E, F)**. To determine whether new OTX2+ neurons resulted from trans-differentiation or MG proliferation, we quantified the number of GFP/OTX2+ cells co-labeled with EdU **(Fig 2G)**. Interestingly, most of the EdU+ infected cells did not express OTX2, (%OTX2+/EdU+GFP+, 18.6% in the ASCL1 condition, **Fig 2G**) showing that most of the infected MG directly trans-differentiated into neurons without re-entering the cell cycle **(Fig 2H)**.

Our success with using the HES1 promoter to stimulate neurogenesis from MG in human 3D cultures, suggests that this promoter could be a more general strategy for neural reprogramming in mice as well. To test whether the HES1 promoter can also drive reprogramming in mouse MG, retinas from P12 and adult mice were cultured as explants and infected with the same lentiviral constructs used for the human. At 7 dpi, retinal explants were fixed and immunolabelled with the same markers used for the retinospheres **(Fig S2D)**. In absence of ASCL1, GFP+ cells expressed the glial markers HES1 and SOX2, validating the specificity of the HES1 promoter for mouse MG **(Fig S2E)**. In the HES1-ASCL1 condition, most GFP+ cells acquired a neuronal morphology and expressed MASH1(ASCL1) and OTX2 **(Fig S2F)**. After ASCL1 expression, we found a similar fraction of GFP/OTX2+ cells as previously observed in human retinospheres (27%) **(Fig S2G)**. In the ASCL1-infected MG, we found some GFP/OTX2+ cells also labelled with EdU demonstrating that they were newborn neurons **(Fig S2H)**. In addition to the P12 mouse retinal explants, we also tested the ASCL1 virus on adult mouse retinal explants; however, we did not detect any GFP/OTX2+ neurons **(Fig S2I, S2J)**. This result is consistent with previous studies demonstrating transgenic expression of ASCL1 alone is not sufficient to stimulate the MG neurogenesis in the adult mouse retina(5,6,24).

### Lentiviral mediated combinations of bHLH increase MG to neuron reprogramming efficiency. (Fig 3) (Fig S3)

Previous studies in mice demonstrated that additional TFs along with ASCL1 influence the cell fate of MG-derived neurons (7,8). To test this strategy in human 3D cultures, we designed an additional lentiviral construct containing ASCL1-ATOH1-GFP under the HES1 promoter. The new construct was first tested on retinospheres, following the same paradigm as previously described **(Fig 3A)**.

**Figure 3:**
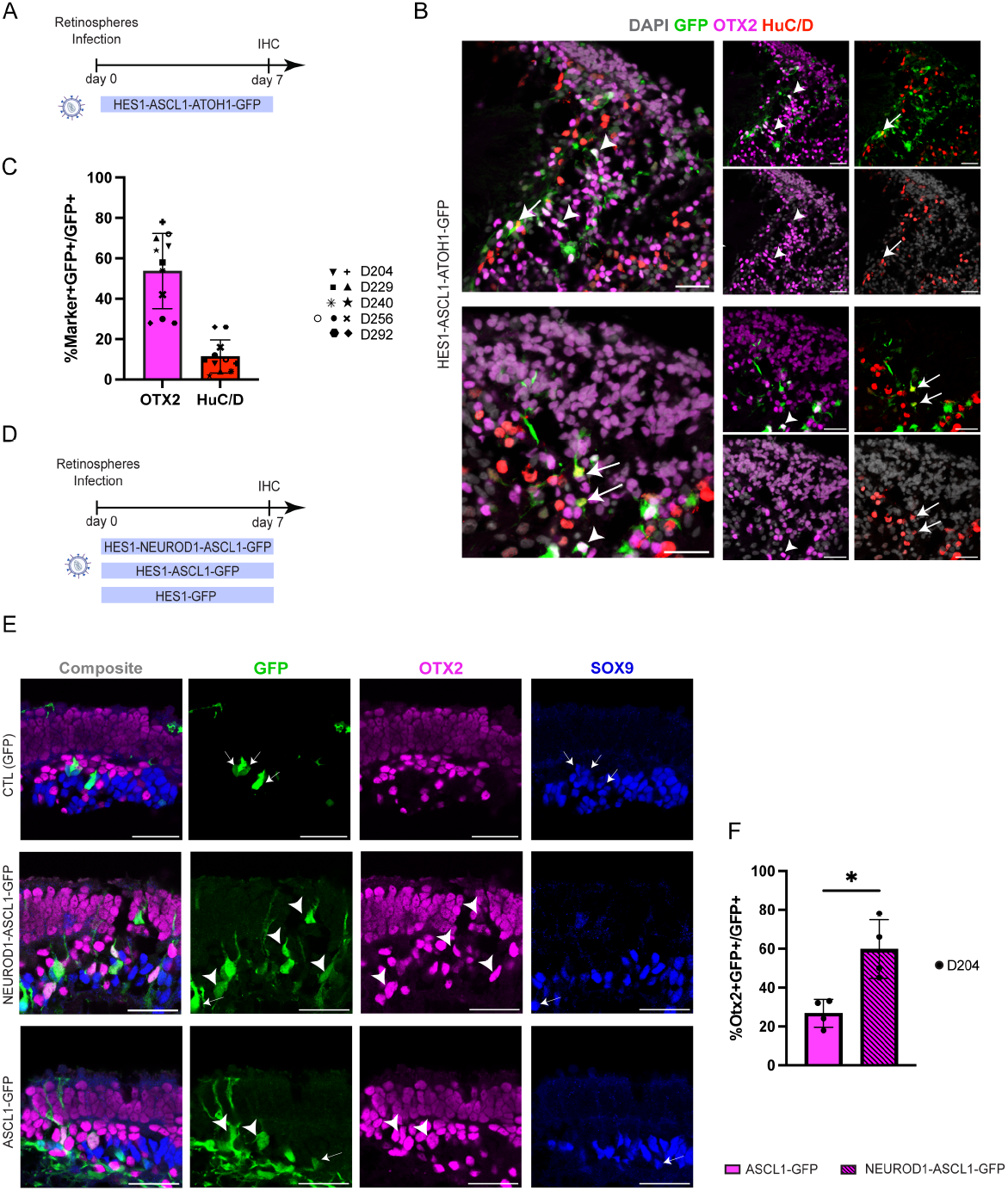
Combo of TFs enhance MG reprogramming efficiency in retinospheres. (A) Schematic of the experimental workflow to test MG reprograming with the HES1-ASCL1-ATOH1-GFP lentivirus. (B) Representative images showing MG-derived neurons expressing OTX2 (magenta) or HuC/D (red) after ASCL1-ATOH1 expression at 7 dpi. Arrowheads show co-labeled GFP/OTX2+ and arrows indicate GFP/HuC/D+ cells. DAPI (grey). Scale bar = 30 *µ*m. (C) Quantification of the MG-derived neurons expressing OTX2 or HuC/D. Each shape represents a retinosphere and ages of the retinospheres are indicated on the right side of the figure. (E) Representative images showing MG-derived neurons expressing OTX2 (magenta) and not SOX9 (blue) 7 days after ASCL1 or NEUROD1-ASCL1 overexpression. Arrowheads show co-labeled GFP/OTX2+ cells. Arrows show GFP/SOX9+ cells. DAPI (grey). Scale bar = 30 *µ*m. (F) Quantification of the MG-derived neurons expressing OTX2. Each dot represents a retinosphere. [8 x D204]. Significance was determined by a Welch’s t-test (*p < 0.05).

We then performed IHC with several neuronal markers. Most of GFP+ cells acquired a neuronal morphology, and we found many GFP+ cells co-localizing with OTX2 or HuC/D **(Fig 3B)**. As OTX2 and HuC/D are markers of distinct populations of neurons (i.e., photoreceptors/bipolar cells and RGCs/amacrine cells, respectively), the results showed that expressing this combination of TFs (ASCL1 and ATOH1) can expand the diversity of neuronal types generated from MG in humans, much like what this combination does in mice(8). In addition, ASCL1-ATOH1 infected cells demonstrated a global increase in reprogramming efficiency compared to ASCL1 alone, similar to our observations in transgenic mice. Interestingly, in human retinospheres, co-expression of ASCL1 and ATOH1 leads to a higher proportion of MG-derived neurons expressing the OTX2 marker compared to HuC/D (53.6% for OTX2 and 11.5% for HuC/D) **(Fig 3C)**. This is different from mice, where the majority of MG-derived neurons expressed HuC/D and acquired an immature RGC phenotype(8).

To determine if this difference between human and mouse MG is maintained using the lentiviral construct, P12 **(Fig S3A, S3B)** and adult mouse **(Fig S3C, S3D)** retinal explants were then infected with the ASCL1-ATOH1 virus. IHC analysis on P12 explants sections **(Fig S3B)** and wholemount **(Fig S3D)** revealed that most GFP+ cells express HuC/D after infection with both ASCL1 and ATOH1 showing that mouse and human MG can respond differently to the same TFs.

As already mentioned, studies in mice demonstrated that combinations of developmentally expressed TFs could affect the reprogramming process(7). We next tested whether combining another developmental pro-neural TF, NEUROD1, along with ASCL1 would enhance MG neurogenesis. We infected retinospheres with HES1-NEUROD1-ASCL1-GFP, HES1-ASCL1-GFP or HES1-GFP lentiviruses **(Fig 3D)**. In the NEUROD1 condition, we observed more GFP+ cells expressing the neuronal marker OTX2 compared to the control condition (60% for NEUROD1-ASCL1 versus 27% for ASCL1 alone) **(Fig 3E, F)**. In addition, we did not observe any GFP/HuCD+ neurons in the NEUROD1 condition.

Since the combination of ASCL1 and NEUROD1 was particularly effective at stimulating neurogenesis in MG from human retina, we also tested this construct on adult mouse retinal explants **(Fig S3E)**. One week post infection, IHC analysis revealed some GFP+ cells that expressed the OTX2 neuronal marker **(Fig S3F)**. Interestingly, the SOX9 glial marker was still expressed in many reprogrammed MG **(Fig S3F)**. Overall, these results demonstrate that combining different TFs along with ASCL1 increased the efficiency of MG reprogramming.

### Combination of ASCL1 and NEUROD1 induces more bipolar-like cells compared to ASCL1 alone. (Fig 4) (Fig S4)

Since both ASCL1 only and NEUROD1-ASCL1 viruses produce MG-derived OTX2+ neurons, we next conducted single-cell RNAseq (sc-RNAseq) to better analyze how these two conditions differed. For this, we infected retinospheres at D203 with HES1-NEUROD1-ASCL1-GFP, HES1-ASCL1-GFP and HES1-GFP **(Fig 4A)**. At 14 dpi, retinospheres were dissociated and the GFP+ cells were isolated using fluorescence-activated cell sorting (FACS) prior processing for scRNA-seq. For all conditions, the GFP+ cell fractions were extracted and processed as individual samples. Datasets were then merged on a single UMAP (uniform manifold approximation and projection) plot **(Fig S4A)** containing different cell clusters that were identified based on the expression of well-known marker genes **(Fig S4B)**. As expected, we found a cluster of MG and a cluster of astrocytes that both express HES1. We also noticed additional cell clusters of cones, rods and amacrine cells, which are present in all samples, and we assume are carried over from the cell sorting **(Fig S4A, S4B)**. We further subset the MG and MG-derived cell clusters for further analysis **(Fig 4B)**. The subset merged UMAP plot contains cell clusters of the following: (1) MG; (2) Proliferating MG; (3) Neurogenic precursors; (4) Bipolar cell precursors; (5) Bipolar cells and (6) Non-identified cells (NDC) **(Fig 4B, 4C)**.

**Figure 4:**
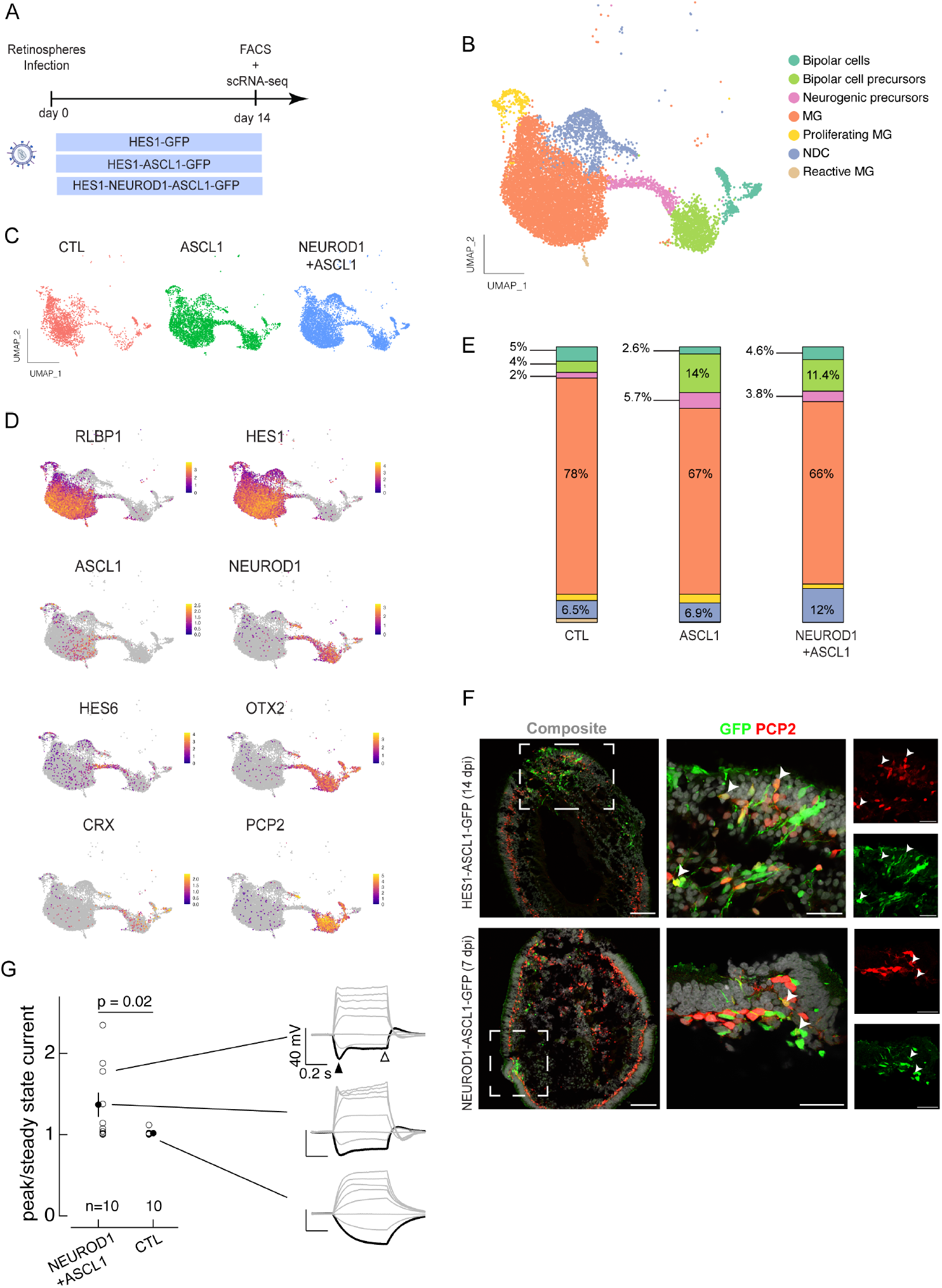
Addition of NEUROD1 along ASCL1 enhances MG reprograming efficiency and pro-mote functional newborn neurons. (A) Schematic of the experimental workflow. (B) Merged UMAP plot of the three conditions (HES1-GFP, HES1-ASCL1, and HES1-NEUROD1-ASCL1). (C) Merged UMAP plot splits by conditions showing the distribution of the different clusters for each condition. (D) Feature plots showing the expression of different genes (*RLBP1, HES1, ASCL1, NEUROD1, HES6, OTX2, CRX* and *PCP2*). (E) Stacked bar plots indicating the percentages of each cell type cluster per condition. (F) Retino-spheres sections labeled with PCP2 (red) and GFP (green). Arrowheads show co-labeled GFP/PCP2+ cells. Scale bar = 100 *µ*m and scale bar = 30 *µ*m in higher magnification. (G) Summary of electrical properties of reprogrammed cells in the NEUROD1-ASCL1 condition 10 days post-transfection. Right three panels show voltage responses to a family of current steps from two reprogrammed cells (top two panels) and one control transfected cell (HES1-GFP) (bottom). The center panel measures the rebound in voltage in response to a hyperpolarizing current step using the ratio of the peak to steady state voltage change (e.g. ratio of voltage changes at the times of the closed and open arrowheads in the top right panel).

Our results confirm the IHC results described above **(Fig 3E)** that ASCL1 and NEUROD1-ASCL1 conditions stimulate the neurogenic capacity of the MG. We found that infection with ASCL1 causes a decrease in expression of RLBP1 and HES1 and an increase in expression of neurogenic precursors genes, such as HES6 **(Fig 4D)**. Furthermore, the addition of NEUROD1 further promotes neurogenic capacity of the MG, since we observed more cells present in the Bipolar cell cluster in this condition compared to ASCL1 alone **(Fig 4D, Fig 4E)**. We detected the expression of both OTX2 and PCP2 in the MG-derived neurons (neurogenic precursors, bipolar cell precursors, and bipolar cells) **(Fig 4D)**. PCP2 is a well-known marker of ON bipolar cell in the mouse retina(25,26). We confirmed the ON-bipolar restricted expression of PCP2 in the fetal human retina using IHC on a D150 retinal section and found PCP2+ cells located in the INL and co-localized with OTX2 **(Fig S4C)**. Next, we used the same PCP2 antibody and confirmed that the protein is expressed in MG-derived neurons in both ASCL1 and NEUROD1-ASCL1 conditions. Interestingly, we were able to detect PCP2 at 7 dpi in the NEUROD1-ASCL1 condition, but only later, at 14 dpi in the ASCL1 condition **(Fig 4F)**.

Since HES1 is also expressed in progenitors, we noticed some residual neurogenesis in the control condition (CTL: HES1-GFP), even without the expression of TFs; this is likely due to the presence of a small pool of progenitors **(Fig 4C)**. To ensure that our results are due to MG reprogramming and not normal developmental neurogenesis, we performed an additional experiment where retinospheres were treated with a notch inhibitor for three days prior to infecting with the viral constructs **(Fig S4D)**; inhibition of notch signaling will force all remaining progenitors to differentiate (27–30). After infection with ASCL1 in the notch treated retinospheres, we found that ASCL1 induced MG-derived OTX2+ cells as well as in the retinospheres that did not receive the notch inhibitor **(Fig S4E)**. Therefore, it is likely that the new neurons we observe in ASCL1-infected retinospheres are arising from MG, not residual progenitors.

Last, we focused on the NEUROD1-ASCL1 condition to test the electrical properties of the MG-derived neurons by performing whole-cell patch clamp recordings from GFP+ cells in retinospheres **(Fig 4G)**. We were particularly interested in whether the complement of ion channels expressed by the cells was affected by TF reprogramming, resulting in changes in the voltage responses to injected current. To test ion channel expression, we measured voltage responses to a family of injected currents. Some of the reprogrammed cells showed a clear rebound in voltage following a hyperpolarizing current step (e.g. black trace in upper right panel of **Fig 4G**). A hyperpolarization-activated current produces similar voltage rebounds in many neurons, including several retinal neuron types. In addition, voltage-clamp recordings revealed a slowly activating inward current following a hyperpolarizing voltage step **(Fig S4F)**. We quantified the rebound in voltage by measuring the ratio of the peak voltage change (filled arrowhead in top right panel of **Fig 4G**) to the steady-state voltage change (open arrowhead). **Figure 4G** plots these ratios for each reprogrammed cell and each matched control cell. Four of ten reprogrammed cells and none of 10 control cells showed such a rebound; on average, the difference in peak vs steady-state ratio was significantly larger in reprogrammed cells compared to matched controls (HES1-GFP). Overall, this result demonstrated that MG-derived cells induced by NEUROD1-ASCL1 display electrophysiological properties different from typical glial cells and more comparable to neurons.

Together our results demonstrated that in both conditions, ASCL1 and NEUROD1-ASCL1, induced MG-derived neurons that resemble bipolar cells. Furthermore, the combination of NEUROD1-ASCL1 stimulates neurogenesis in MG with a higher efficiency than ASCL1 alone.

### MG reprogramming in the human adult postmortem retina. (Fig 5)

The results with the fetal retinospheres demonstrated that human MG can be reprogrammed to neural progenitors to regenerate retinal neurons; however, it is not known whether adult human MG can be reprogrammed to neural progenitors. This is a critical next step to evaluate the feasibility of in vivo reprogramming for regenerative medicine. To determine whether neurogenesis can be stimulated in adult human MG, we used human adult postmortem retinas, cultured as explants. Previous studies have shown that forced expression of ASCL1 alone in mature mouse MG is no longer sufficient to induce MG reprogramming(5,6). However, the combination of ATOH1 along with ASCL1 has proven to be efficient in reprogramming adult MG in the mouse retina(7,8). Therefore, we tested several combinations of reprogramming factors in adult human post-mortem retinas.

**Figure 5:**
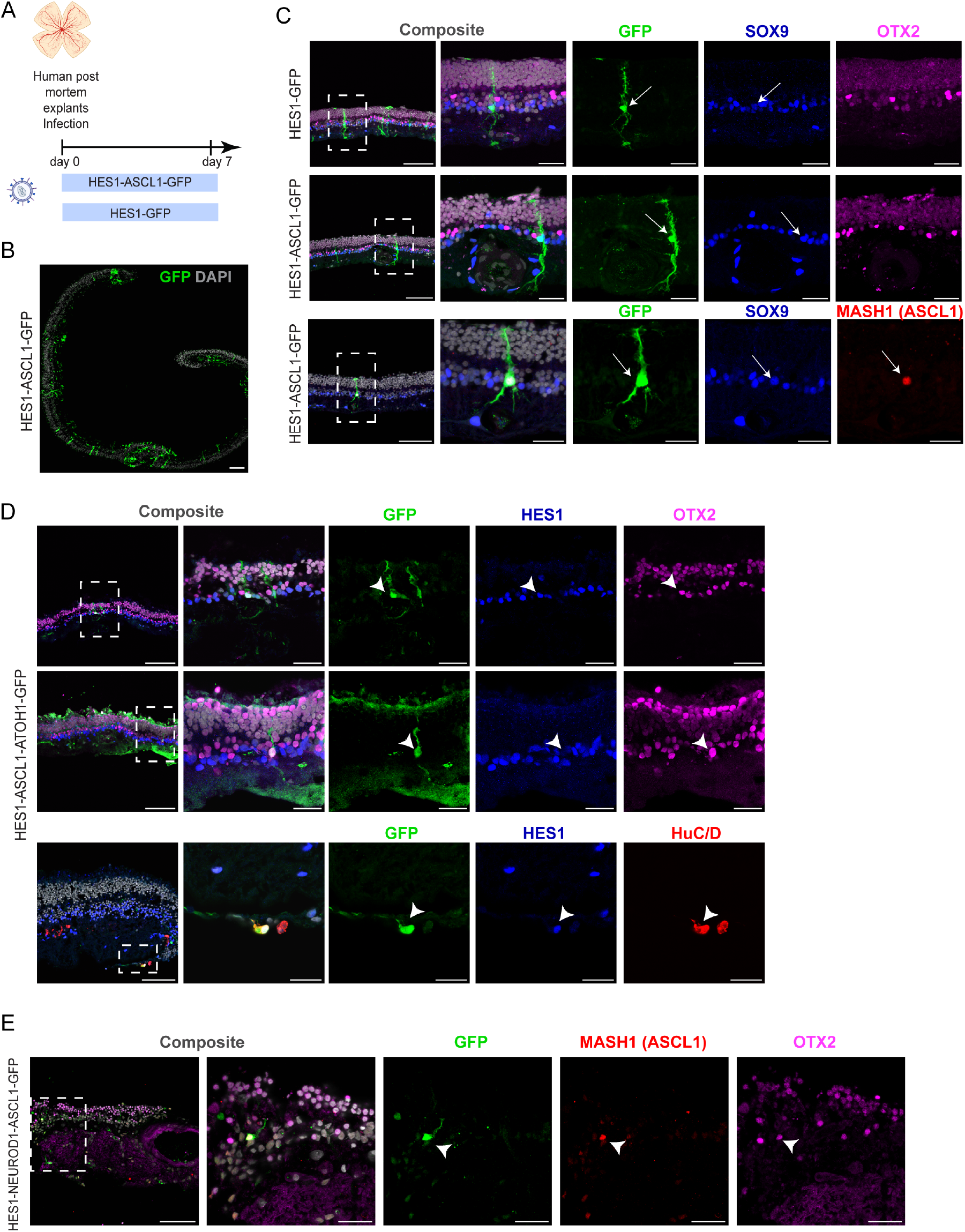
MG reprogramming in the adult human retina *in vitro*. (A) Schematic of the experimental workflow to test MG reprograming in adult post-mortem retinal explants. (B) Representative images showing the infection efficiency with the HES1-ASCL1 lentivirus in the adult post-mortem retinal explants. DAPI (grey) and GFP (green). Scale bar = 100 *µ*m. (C) Representative images showing that HES1 drives the expression of ASCL1 (MASH1, red), but it is not sufficient to reprogram human adult MG in vitro. SOX9 (blue), GFP (green), OTX2 (magenta), MASH1 (ASCL1, red), and DAPI (grey). Arrows indicate GFP/SOX9+ or GFP/SOX9/MASH1+ cells. Scale bar = 100 *µ*m and scale bar = 30 *µ*m in higher magnification. (D) Example images of MG-derived neurons after ASCL1-ATOH1 expression. HES1 (blue), GFP (green), OTX2 (magenta) and DAPI (grey). Arrowheads show co-labeled GFP/HES1/OTX2+ or GFP/HES1/HuC/D+ cells. Scale bar = 100 *µ*m and scale bar = 30 *µ*m in higher magnification. (E) Example images of MG-derived OTX2+ neurons after NEUROD1-ASCL1 expression. GFP (green), OTX2 (magenta), MASH1 (ASCL1, red), and DAPI (grey). Arrowheads indicate GFP+ cell co-labeled with MASH1 and OTX2. Scale bar = 100 *µ*m and scale bar = 30 *µ*m in higher magnification.

Retinal explants were initially infected with HES1-GFP or HES1-ASCL1-GFP lentivirus and fixed 7 dpi prior to immunolabelling with different glial and neuronal markers **(Fig 5A)**. We observed a widespread lentiviral infection on human post-mortem explants **(Fig 5B)**. We first confirmed the specificity of the HES1 promoter; all GFP+ expressing cells were colabeled with the SOX9 glial marker (%SOX9+GFP+/GFP+, 100%). In the control condition, cells retained a MG morphology, with processes spanning the entire retina **(Fig 5C)**. Of note, some explants also contained some GFP/SOX9+ cells in the ganglion cell layer (GCL), showing that astrocytes were also infected. In the ASCL1 condition, despite ASCL1 expression (GFP/MASH1+cells), GFP+ cells still retained a MG morphology and did not express OTX2 **(Fig 5C)**. Therefore, ASCL1 alone was not sufficient to stimulate neurogenesis from adult human MG, though it had this potential in the fetal MG.

Results from mice demonstrated that the combination of ASCL1 and ATOH1 was much more efficient in stimulating neurogenesis in adult MG. Therefore, we next infected adult postmortem retinal explants with a virus expressing both ASCL1 and ATOH1, following the same paradigm as described above. Due to the size of the polycistronic transgene cassette, where the fluorescent reporter is on the third position, we detected fewer GFP+ cells in this condition. However, some GFP+ cells exhibited a neuronal-like morphology, with a rounder and smaller nucleus, contrasting with the two previous conditions (HES1-GFP and HES1-ASCL1-GFP). Further IHC analysis showed GFP+ cells expressed OTX2 **(Fig 5D)**. In rare case, we also found some GFP/HuC/D+ cells **(Fig 5D)**. In both cases, the glial/progenitor marker HES1 was still detectable in the MG-derived neurons and not observed in other conditions (HES1-GFP and HES1-ASCL1). Similarly, we also found some GFP/OTX2/MASH1+ cells in the HES1-NEUROD1-ASCL1 condition **(Fig 5E)**. Therefore, the labelling of HES1 or MASH1 (ASCL1) with neuronal markers such as OTX2 and HuC/D further validated that neurons are derived from MG, since this labeling was not present in the control. Taken together, our results demonstrate that MG can be reprogrammed into neurons in the adult human retina using a combination of proneural bHLH TFs.

## Discussion

In the last ten years, studies in mouse, and more recently human, have shown that MG can be stimulated to generate neurons (6–9,18,31). In this study, we aimed to determine whether this same approach can be used in human retina: can human MG be reprogrammed to a neurogenic state. These studies are of necessity in vitro, and so we employed explant cultures of both adult human postmortem retina and human fetal retina (18,32,33). We identified a promoter from the HES1 gene that is active in both glia and progenitor cells, to drive the expression of pro-neural factors, including ASCL1, ATOH1 and NEUROD1. The HES1 promoter has the unique advantage of showing glial specificity to initiate the reprogramming process, as well as expression in the glial-derived progenitor cells, thus providing a novel approach to design of reprogramming strategies. We used this novel design to show that human MG can be stimulated to generate neurons in both adult and fetal retinas. This provides a first proof of concept that in vivo reprogramming to stimulate neurogenesis from glia can provide a fundamentally new approach for repairing the CNS in humans.

The expression of the proneural TF Ascl1 can stimulate neurogenesis from MG in mice, and now we demonstrate that this is possible in human glia as well: MG-specific expression of ASCL1 alone in the retinosphere MG produces OTX2+ neurons. Although the MG reprogramming efficiency can be greater than 20% in retinospheres with ASCL1, we noticed a decrease in MG neurogenesis with the age of the retinospheres and there were no MG-derived neurons in the adult retina after ASCL1 overexpression alone. In younger retinospheres, the higher reprogramming efficiency could partially be explained by the presence of residual progenitors (or less mature MG), particularly in retinospheres from the most peripheral regions of the retina. Furthermore, we observed a low level of neurogenesis in the GFP control condition in our scRNA-seq data. In future studies, it will be important to understand the role of glial maturation in their ability to become neurogenic. Nonetheless, the difference in reprogramming efficiency between fetal and adult human MG mirrors that observed in mice, with a progressive loss of the MG neurogenic capacity over time due to epigenetic changes(6,24). In the adult mouse retina, this issue has been overcome using Trichostatin A (TSA), an epigenetic modulator, along with expression of ASCL1(6–8).

Although the expression of ASCL1 alone was not sufficient to stimulate neurogenesis in adult human MG, we have previously shown that combinations of proneural TFs can substantially increase the efficiency of the reprogramming process in mice. Therefore, we also tested combinations of proneural TFs in the human retinal explants and found that combining ASCL1 with either ATOH1 or NEUROD1, significantly increases the efficiency of neurogenesis in fetal MG. The combination of ASCL1 and NEUROD1 increases the rate of reprogramming from 20% with ASCL1 alone, to over 60%. The increase in reprogramming efficiency further enables the reprogramming of adult postmortem retinal MG; with ASCL1 alone we did not observe neurogenesis in adult retina, but with the combination of ASCL1 with either ATOH1 or NEUROD1, we found many examples of MG-derived neurons of either bipolar or amacrine/RGC fate.

We have also found in mice that different combinations of reprogramming factors direct the progeny of MG to specific retinal cell fates: over-expression of ASCL1 induces MG to generate primarily bipolar cells, while most MG-derived neurons expressed HuC/D after overexpression of ASCL1 and ATOH1(5). Although we find similar results with the human MG, there are some differences. For example, while nearly 80% of the neurons generated from MG over-expressing Ascl1 and Atoh1 resemble ganglion cells, a similar combination of TFs in human retina resulted in fewer than 20% of the MG-derived neurons expressing HuC/D. It is not clear why human and mouse MG would behave differently with the same combination of TFs and the identical delivery system. However, this discrepancy between species highlights the importance of testing combinations of already known TFs in a human model, as well as exploring additional TFs to potentially unlock still inaccessible cell fates.

Nonetheless, an important advance in our findings is that combining TFs along with ASCL1 can reprogram MG in the adult human retina without the addition of epigenetic modulators. This represents a proof of concept that we can stimulate neurogenesis from human MG using a viral vector. For future clinical applications, it will probably be necessary to opt for a non-integrative viral vector strategy such as AAVs (adeno-associated virus), though it will be important to control for the fact that the specificity of the glial promoter GFAP, when delivered via an AAV vector, changes over time when used to express pro neural TFs (23–25,41).

Overall, this study constitutes a critical step that will advance in vivo MG reprogramming as a regenerative approach for the retina. Although many barriers still need to be overcome, the fact that neurogenesis can be stimulated in adult human MG shows that this approach may someday lead to a therapeutic strategy to treat patients suffering from retinal diseases.

## Acknowledgements

We would like to thank Dr. Ian Glass and the members of the BDRL (Birth Defect Research Laboratory) for their valuable help with the human fetal tissues. Thank you to all the members of the Reh lab (past and present) and the Bermingham-McDonogh lab for their valuable comments on the manuscript.

This work is funded by a grant from the Foundation Fighting Blindness (TA-RM-0620-0788-UWA) to T.A.R. and by a sponsored research agreement with Tenpoint Therapeutics.

## Author contributions

Overall conceptualization, J.W. and T.A.R.; retinosphere cell culture, mouse retinal explants and cell infection, J.W., F.K., retinosphere and mouse retinal explant tissue processing, J.W., F.K., D.H., and K.A.; Human adult tissue acquisition and infection, G.L., N.S. and A.M.; Immunohistochemistry and microscopy, J.W., F.K. D.H., and K.A.; single cell genomics data analysis: J.W., and T.A.R.; electrophysiology experiments, F.R.; manuscript preparation and writing, J.W. and T.A.R.; funding acquisition, T.A.R.; and supervision and project administration, T.A.R.

## Supplemental information

**Document S1**. Figures S1–S4.

**Table S1:** List of the viral vectors.

**Table S2:** List of antibodies used for IHC.

## Materials and Methods

### Retinospheres culture

Human retinal tissues were provided from the Birth Defect Research Laboratory at the University of Washington following the approved protocol Retinospheres (UW5R24HD000836). Retinospheres were cultured as previously debrided (18,32). Briefly, retinospheres are maintained in low binding attachment plate in Retinal Differentiation Medium (RDM 5%; supplemented with 5% fetal bovine serum (FBS), DMEM/DMEM/F12, 2% B27, 1% Pen-Strep) at 37ºC incubator with 5% CO2.

### Postmortem Human and mouse retinal explants culture

De-identified cadaveric posterior eye sections were obtained from Sierra Donor Services Eye Bank (West Sacramento, CA) for this study. These were research grade tissues deemed unsuitable for transplantation, with consent obtained for their use in research in accordance with the Uniform Anatomical Gift Act, Ca. Health and Saf. Code § 7150, Eye Bank Association of America Medical Standards, and Sierra Donor Services Eye Bank Standard Operating Procedures. The peripheral retina, excluding the macula, was meticulously dissected and washed in HBSS containing antibiotics and then carefully segmented for retinal explant culture. Retinal segments approximately 3 mm square were placed with the photoreceptor side facing up onto a 0.4 *µ*m pore culture insert in a 6 well plate. Medium (Neural Basal medium containing 10% FBS, 1% N2, 1% B27, 0.5% L-Glutamine and 1% PenStrep) was changed every other day. Similarly, the same protocol was followed for P12 or adult mouse retinal explants. Virus (Table 1) was pipetted directly onto the surface of the explant. Retinal explants were maintained in culture in incubator at 37ºC with 5% CO2.

### Vector transduction in 3D Retinospheres

Retinospheres of desired ages were collected and dissected in half. Each retinospheres was then placed in an individual well of a normal attachment 96 well plate. For each condition, individual retinosphere received the same master mix containing RDM5%, Neuraminidase (MilliporeSigma), Polybrene, DMSO, EdU and the virus (Table 1). The following day, retinospheres were transferred to an individual well of a low binding attachment plate without removing the medium. The next day the medium was replaced with fresh RDM5% and EdU. Viral vectors were sources from VectorBuilder (Table 1).

### Tissue preservation and IHC

Tissue preservation and immunostaining were performed as previously described (18). Briefly, Retinal Tissues (Retinospheres and retinal explants) were fixed with PFA4% for 15 min prior to embedding in successive solutions containing 10%, 20%, or 30% sucrose for 30 mins each at RT. Tissues were then frozen in OCT and cryosectionned at 15 *µ*m for retinospheres and 18 *µ*m for retinal explants (mouse and human). For immunostaining, sections were washed three times in PBS and then incubated with a blocking solution (10% horse serum, 90% PBS, and 0.5% Triton X-100) for 1h at RT. Primary antibodies (Table 2) were diluted in the blocking solution and incubated overnight at 4ºC. The following day, sections were washed three times with PBS prior to a 1h treatment at RT with secondary antibodies (Table 2) primarily diluted in blocking solution containing 1/8000 DAPI. After three PBS washes, sections were coverslipped using Fluoromount-G (SouthernBiotech) mounting medium. For EdU labeling, the Click-it solution (Click-iT EdU Assay, Invitrogen) was used following the manufacturer’s instructions.

### Microscopy/cell counts and statistical analysis

Imaging was performed using a Zeiss LSM880 confocal microscope. For quantification analysis with cell type specific markers (OTX2, HuC/D, SOX9, HES1), a total of 50 GFP+ cells per retinospheres were included from random sections. GFP/Marker+ cells were then counted with a 20X or 63X objective. Each dot/shape on the bar graph represents an individual retinosphere. Graphics and statistical analysis were performed using GraphPad Prism (version 10.2.2). Welch’s t-test were used for analysis between two conditions. Results are presented as mean ± SD, except **Fig 2G**, results are presented as mean ± SEM.

### Electrophysiology

We recorded electrical responses of transfected GFP+ cells using whole-cell patch clamp techniques. Fluorescent cells were identified using a two-photon microscope, and GFP expression in all recorded cells was confirmed by filling the cells with a second dye (Alexa 594) during recording and checking colocalization with GFP. Patch pipettes were filled with an internal solution containing 123 mM K-aspartate, 10 mM KCl, 10 mM HEPES, 1 mM MgCl2, 1 mM CaCl2, 2 mM EGTA, 4 mM Mg-ATP, 0.5 mM Tris-GTP, and 100 *µ*m Alexa Fluor 594-hydrazide. Whole cell patch pipettes had resistances of 12-14 MΩ. Access resistance was < 25 MΩ for all cells. Reported voltages have not been corrected for a ∼-10 mV liquid junction potential.

### Fluorescent Activated Cell Sorting (FACS)

Retinospheres were dissociated into single cells as previously described(18). Following dissociation cells were passed through a 35 *µ*m strainer. Fluorescent Activated Cell Sorting (FACS) was performed to isolate the GFP+ cells using a BD FACSAria III cell sorter (BD Bioscience). Single cell RNA library construction

Following FACS purification, GFP+ cells were centrifuged at 300g at 4ºC for 5 mins. For each sample, cells were resuspended in RDM5% to reach a concentration of 1000 cells per µl. Libraries construction were done using the Chromium Next GEM single Cell 3’ Reagent kits v3.1 (Dual Index) following the manufacturer’s instructions. Libraries were next sequenced using Illumina NextSeq.

### Data code and availability

All Single cell data from this study is available on as GEO XXXX

## Supplementary Information

**Table S1:**
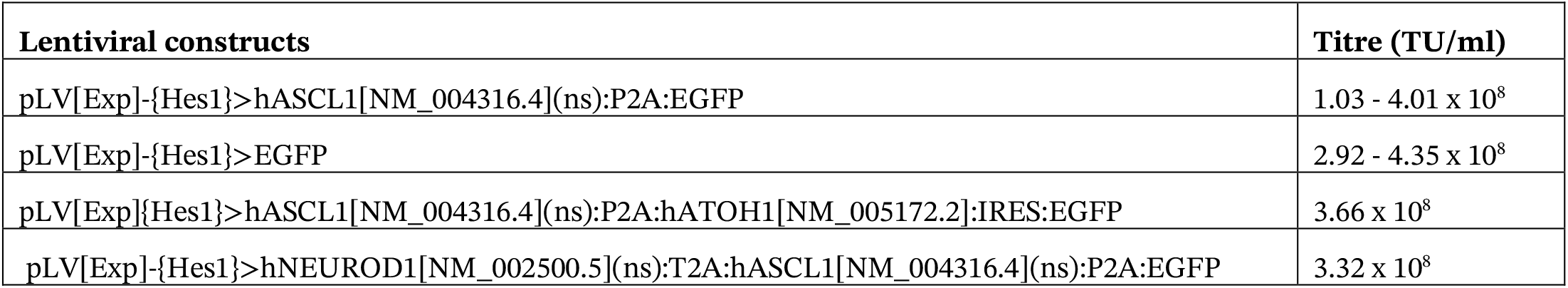
List of the viral vectors.

**Table S2:** List of antibodies used for IHC.

| Antibodies | Source (brand) | Identifier (catalogue identifier) | Dilution |
| --- | --- | --- | --- |
| Rabbit anti-SOX9 | Millipore | Ab5535 | 1/300 |
| Goat anti-OTX2 | R&D Systems | BAF1979 | 1/300 |
| Mouse anti-HUCD | Invitrogen | A-21271 | 1/100 |
| Chicken anti-GFP | abcam | Ab13970 | 1/300 |
| Rabbit anti-Hes1 | Cell Signaling | D6P2U | 1/300 |
| Rabbit anti-MASH1 | abcam | Ab211327 | 1/300 |
| Mouse anti-PCP2 | Santa Cruz Biotechnology | sc-137063 | 1/300 |
| Rabbit anti-VSX1 | proteintech | 23566-1-AP | 1/300 |
| Donkey anti-mouse 647 | Invitrogen | A32787 | 1/300 |
| Donkey anti-rabbit 647 | Invitrogen | A31573 | 1/300 |
| Donkey anti-mouse 568 | Invitrogen | A10037 | 1/300 |
| Donkey anti-goat 568 | Invitrogen | A11057 | 1/300 |
| Donkey anti-chicken 488 | Jackson ImmunoResearch | 703-546-155 | 1/300 |

**Figure S1:**
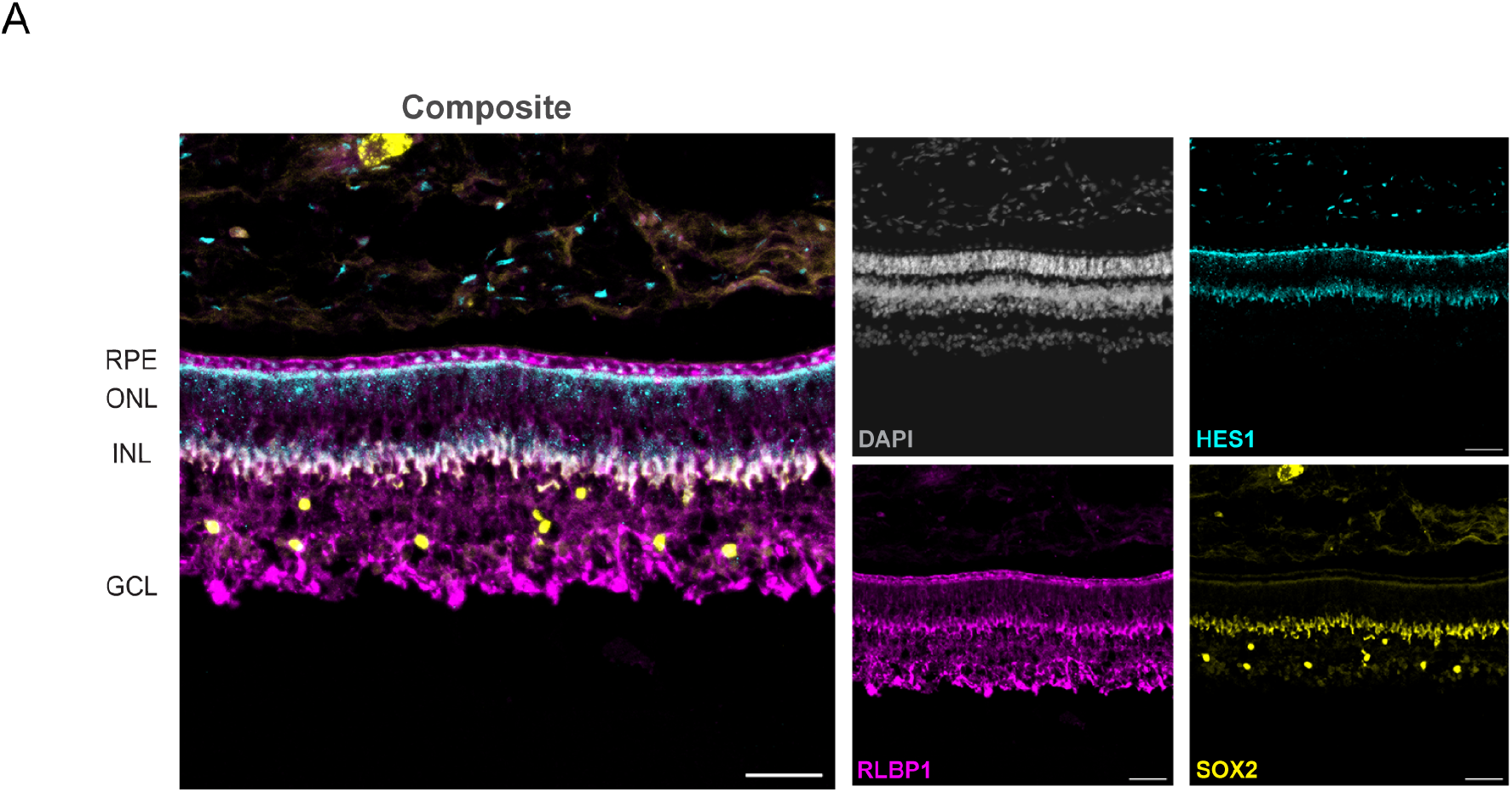
HES1 is expressed in MG in the human fetal retina. (A) Representative images showing expression of HES1 in a fetal retinal section at 150 days. HES1 (cyan), SOX2 (yellow), RLBP1 (magenta), and DAPI (grey). Scale bar = 50 *µ*m.

**Figure S2:**
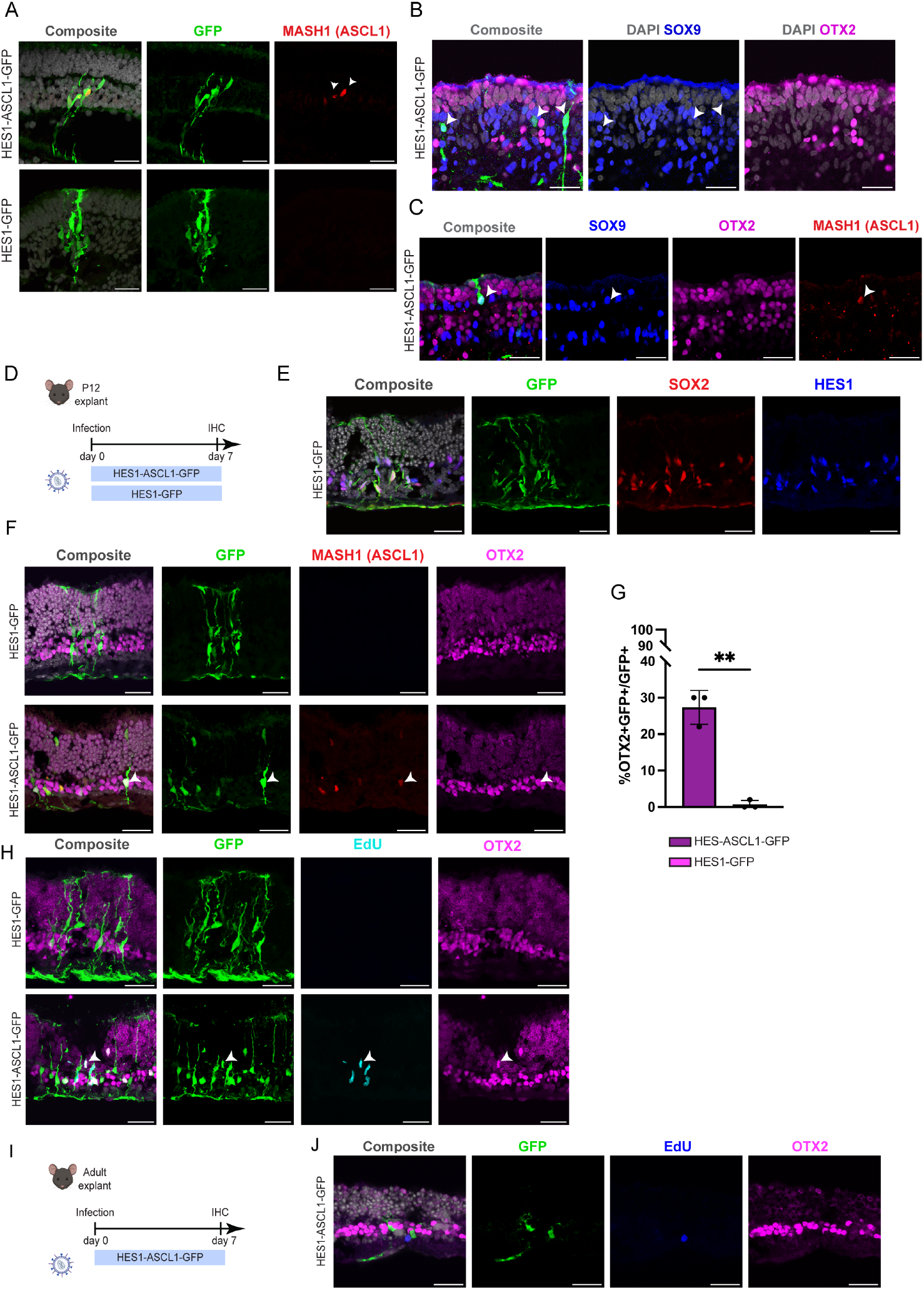
Specificity of the HES1-ASCL1-GFP lentiviral vector in retinospheres and mouse MG reprogramming *in vitro*. (A) Representative images of retinospheres infected with HES1-GFP or HES1-ASCL1-GFP and immunolabeled for GFP (green), DAPI (grey) and ASCL1 (MASH1, red). Arrowheads show co-labeled GFP/MASH1+ cells. Scale bar = 30 *µ*m. (B), (C) Representative images immunolabeled for GFP (green), DAPI (grey) SOX9 (blue), and OTX2 (magenta), 48 hours post infection with HES1-ASCL1-GFP. Arrowheads indicate co-labeled GFP/SOX9+ or GFP/SOX9/MASH1+ cells. Scale bar = 30 *µ*m. (D) Schematic of the experimental workflow to test MG reprograming in P12 mouse retinal explants. (E) Representative images showing the glial specificity of the HES1 promoter in mouse retinal explants. GFP (green), SOX2 (red), HES1 (blue), and DAPI (grey). Scale bar = 30 *µ*m. (F) Representative images of MG-derived OTX2+ cells after ASCL1 expression in mouse retinal explants. GFP (green), ASCL1 (MASH1, red), OTX2 (magenta), and DAPI (grey). Arrowheads indicate GFP/ASCL1/OTX2+ cells. Scale bar = 30 *µ*m. (G) Quantification of the percent of GFP+ MG-derived neurons that express OTX2, 7-dpi. Each data point was calculated from 50 GFP+ cells from n = 3 mice. Scale bar = 30 *µ*m. Significance was determined by a Welch’s t-test (**p < 0.01). (H) Confocal images showing EdU/OTX2+ MG-derived neurons after ASCL1 overexpression in P12 mouse retinal explant. GFP (green), EdU (cyan), OTX2 (magenta). Arrowheads indicate GFP/EdU/OTX2+ cells. Scale bar = 30 *µ*m. (I) Schematic of the experimental workflow to test MG reprograming on adult mouse retinal explants with the HES1 -ASCL1-GFP lentivirus. (J) Representative image showing ASCL1 overexpression is not sufficient to reprogram adult mouse MG in OTX2+ neurons. GFP (green), OTX2 (magenta), EdU (blue), and DAPI (grey). Scale bar = 30 *µ*m.

**Figure S3:**
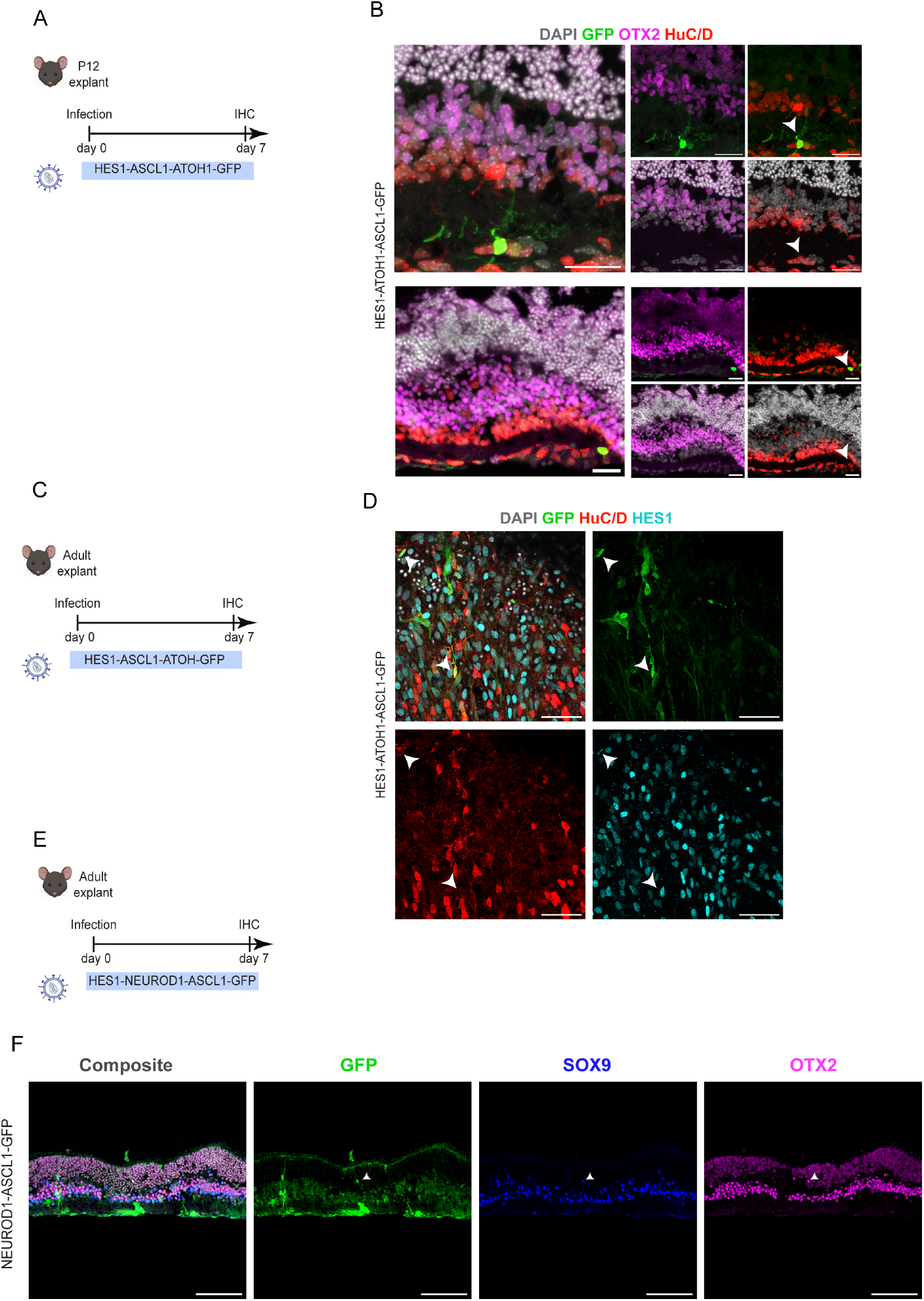
Combination of ASCL1 and ATOH1 or NEUROD1 can reprogram adult mouse MG *in vitro*. (A) Experimental workflow to test HES1-ASCL1-ATOH1 lentivirus on P12 mouse retinal explants. (B) Example images showing MG-derived HuC/D+ neurons after ASCL1-ATOH1 overexpression. GFP (green), OTX2 (magenta), HuC/D (red), and DAPI (grey). Scale bar = 30 *µ*m. Arrowheads indicate GFP/HuC/D+ cells. Scale bar = 50 *µ*m. (C) Experimental workflow to test HES1-ASCL1-ATOH1 lentivirus on adult mouse retinal wholemount. (D) Wholemount images showing MG-derived HuC/D+ neurons after ASCL1-ATOH1 overexpression. GFP (green), OTX2 (magenta), HuC/D (red), and HES1 (cyan). Scale bar = 30 *µ*m. Arrowheads indicate GFP/HuC/D+ cells. Scale bar = 30 *µ*m. (E) Schematic of the experimental workflow to test MG reprograming on adult mouse retinal explants with the HES1-NEUROD1-ASCL1 lentivirus. (F) Example images showing MG-derived OTX2+ cells. GFP (green), SOX9 (blue), OTX2 (magenta), and DAPI (grey). Arrowheads show co-labeled GFP/SOX9/OTX2+ cells. Scale bar = 100 *µ*m.

**Figure S4:**
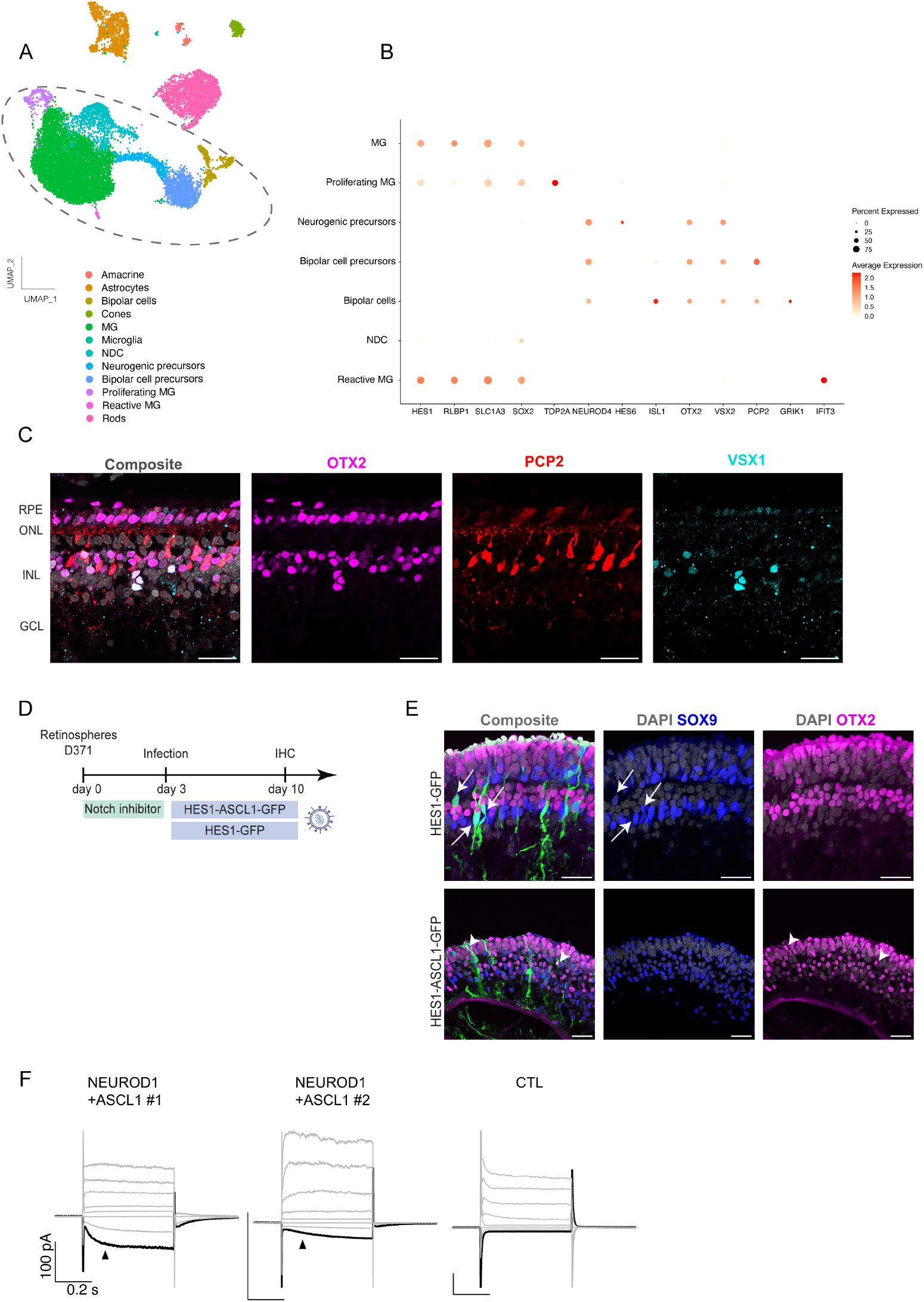
Further characterization of neurogenesis from human fetal MG. (A) Combined UMAP plot of the three different conditions (HES1-GFP, HES1-ASCL1, and HES1-NEUROD1-ASCL1-GFP) with the cell clusters identified by the expression of well-known marker genes. Dashed circle represents the subset clusters present in Figure 4B. (B) Dot plot showing the expression of marker genes for the different cell clusters composing the UMAP plot. (C) Representative images of a fetal retinal section at 150 days showing PCP2 is expressed in bipolar cells. OTX2 (magenta), PCP2 (red), VSX1 (cyan), and DAPI (grey). Scale bar = 30 *µ*m. RPE, retinal pigmented epithelium; ONL, outer nuclear layer; INL, inner nuclear layer; GCL, ganglion cell layer. (D) Schematic of the experimental workflow. A notch inhibitor was added to the retinospheres prior to infecting with HES1-ASCL1-GFP or HES1-GFP. (E) Representative images showing MG-derived OTX2+ neurons after ASCL1 overexpression in retinospheres that previously treated with a notch inhibitor (gamma secretase inhibitor (PF4014)). OTX2 (magenta), SOX9 (blue), and DAPI (grey). Arrows indicate GFP/SOX9+ cells and arrowheads show GFP/OTX2+ cells. Scale bar = 30 *µ*m. (F) Examples of voltage-clamp recordings of MG-derived neurons after HES1-NEUROD1-ASCL1-GFP or HES1-GFP infection (CTL).

## Notes

### Competing Interest Statement

This research is funded in part by a sponsored research agreement with Tenpoint Therapeutics; T.A.R. is a co-founder and consultant. Some of the findings in this manuscript are part of a patent that has been submitted by the University of Washington: PCT/US23/65219.

